# Development of *Encarsia tabacivora* (Viggiani) (Hymenoptera: Aphelinidae) in nymphs of *Bemisia tabaci* (Gennadius) MEAN 1 (Hemiptera: Aleyrodidae)

**DOI:** 10.1101/2023.01.06.486630

**Authors:** Heidy Gamarra, Marc Sporleder, Luz Supanta, Alexander Rodríguez, Jürgen Kroschel, Jan Kreuze

## Abstract

*Encarsia tabacivora* Viggiani 1985 (Hymenoptera: Aphelinidae), is an endoparasitoid of whiteflies, including *Bemisia tabaci* (Gennadius) MEAN 1 (Hemiptera: Aleyrodidae), reported from southern USA, Caribbean, and Brazil. In field surveys, the natural occurrence of *E. tabacivora* was confirmed in the Cañete valley of Peru. Its biology and development was studied under laboratory conditions at 20°C and 70-75% RH. The results showed that fertilized females lay their eggs inside nymphs of the 3^rd^ and early 4^th^ instar of *B. tabaci* MEAN 1. The stages of development are egg, three larval instars, prepupa and pupa. Development from egg to adult lasted 19.3 (SE±0.17) days for females. No males were produced, which indicates that *E. tabacivora* exhibits thelytokous parthenogenesis. Parasitized host nymphs exhibited multiple oviposition punctures and several L_1_-instar wasps were observed in individual host nymphs, indicating superparasitism. However, from the L_2_-instar onwards only one individual developed to adult stage. *E. tabacivora* might be a prospective biological control agent for *B. tabaci* in sweet potato also for minimizing the spread of whitefly-induced plant viruses in Peru and other regions of the world. The knowledge gained might be helpful in establishing mass rearing protocols of *E. tabacivora* for inoculative releases.

## 1 Introduction

The Encarsia Föster genus is one of the most diverse and economically important group of parasitoids among the family Aphelinidae, which are used for biological control. This genus contains more than 437 valid species (Polaszek et al. 2004, Noyes 2015) of which about 150 species are used globally in classical biological control programs against whiteflies (Aleyrodidae) and armored scale insects (Diaspididae) in a broad range of agricultural and vegetable crops (Myartseva and Evans 2007).

De Santis (1981) described the species *Encarsia tabacivora* as *E. bemisiae*, however, since the name was homonymously already used for describing another species, Viggiani (1985) changed the name to *E. tabacivora*. Polaszek et al. (1992) considered *E. tabacivora* Viggiani as a synonym for *E. pergandiella* Howard, although yet unpublished studies by J. B. Woolley and R. Johnson strongly suggest *E. tabacivora* as a valid species (Hernández-Suárez et al. 2003). More recent research confirmed that *E. tabacivora* and *E. pergandiella* are different species based on characters such as the number of setae on the sixth tergite, the relative length of the ovipositor to midtibia, and of antennal segments (Myartseva and Evans 2007); albeit the authors stated that both species have a light (pupating face up) and a dark form (pupating face down). Both species *E. tabacivora* and *E. pergandiella* are important biological control agents of whiteflies (Hemiptera: Aleyrodidae), like the silverleaf whitefly *Bemisia tabaci* (Gennadius) (sensu lato), and the greenhouse whitefly, *Trialeurodes vaporariorum* (Westwood) (Gebiola et al. 2017).

*E. pergandiella* appears to have a wide geographic distribution, being present in the Americas (Myartseva and Evans 2007), as well as in the Old World, although only the dark form was observed (Viggiani 1988, Hernández-Suárez et al. 2003). By contrast, *E. tabacivora* has only been recorded from the Americas. *E. tabacivora* has been reported from Brazil (Gebiola et al. 2017), Caribbean (including West Indies and the Dominican Republic) (Evans and Serra 2002), Mexico (Myartseva and Evans 2007), and some states of the United States of America (Arizona, California, South Carolina, and Texas) (Gebiola et al. 2017).

In Peru, *E. pergandiella* has been released in 2001 as a biocontrol agent for whiteflies in the La Libertad region. Under greenhouse conditions, a parasitism level of 85% was found; however, in open fields, the level of parasitism was low which was explained by the use of insecticides to control other pests (Díaz and Ternero 2002). There is, however, no report which confirms the permanent establishment of *E. pergandiella* in Peru.

In this study, parasitoids of the whitefly *B. tabaci* were collected south of Lima in the Cañete valley from cassava (*Manihot esculenta* Crantz) and sweet potato (*Ipomoea batatas* (L.) LAM.) fields free of chemical management. This preliminary survey revealed that naturally occurring parasitoids of whiteflies could only be detected from insecticide-free farmers’ fields, while fields that were regularly treated with insecticides were parasitoid-free. This indicated that the widespread use of insecticides attributed to a significant reduction of the abundance of parasitoids in these crops. The collected specimens of *Encarsia* were identified as *E. tabacivora* and *E. sophia* (Girault & Dodd 1915) (by specialists at APHIS, USDA, USA), assumed to be endemic species in the study area and reported for the first time to occur in Peru (Supanta 2017). Subsequently, the two species were reared under controlled conditions in the laboratory, whereby *E. tabacivora* was easier to rear also having higher parasitism rates in *B. tabaci* than *E. sophia*.

Therefore, further studies, including the present study, have been initiated on *E. tabacivora* as a prospective indigenous agent for biological control of whiteflies in Peru.

The reproduction of aphelinids is commonly biparental, although uniparental reproduction also occurs frequently. Usually, Encarsia species, as other hymenopterans, have a haplo-diploid sex determination system and reproduce by arrhenotokous parthogenesis, that is, unmated females lay diploid eggs in or on their host’s body that invariably give rise to females, whereas males develop from haploid, unfertilized eggs by arrhenotokous parthenogenesis (Myartseva et al. 2012). Although arrhenotokous parthogenesis is usually the case in Encarsia species, in rare cases thelytokous parthogenesis, as for example in *E. formosa* Gahan, have been found (i.e., diploid females develop from unfertilized eggs) with rare or unknown males. All thelytokous species of Encarsia examined to date have been found to be infected by parthenogenesis-inducing endosymbiotic bacteria, e.g., of the genus Wolbachia in *E. formosa* (Zchori-Fein et al. 2001) and of the genus Cardinium in all other examined species (Zchori-Fein et al. 2001, 2004, Giorgini et al. 2009, Gebiola et al. 2017). It has been shown that populations from different geographic regions have specific reproductive modes depending on their endosymbiont infection status. The *E. tabacivora* population from Brazil, for example, reproduces exclusively by thelytokous parthenogenesis and is infected by a strain (cEper2) of Cardinium (Zchori-Fein et al. 2001, 2004, Gebiola et al. 2017).

Currently, because of global warming, it is very likely that the sweet potato whitefly *B. tabaci* MEAN 1 may expand its distribution to new geographical regions as well as increase its abundance in regions where it already exists, being a potential risk for a wide range of host crops (Gamarra et al. 2016). Since in Peru *B. tabaci* MEAN1 is a significant pest in sweet potato that is difficult to control due to its evolved high resistance against chemical pesticides including broadly used growth regulators like buprefezin, alternative pest management strategies need to be developed, using naturally occurring antagonists as much as possible. Antagonists might play an important role in preventing crop losses in agroecosystems and their use as biological control agents should be thoroughly evaluated. As a newly identified naturally-occurring antagonist of *B. tabaci* in Peru, studies on the biology and potential use of *E. tabacivora* have been initiated. The objective of the present study was to determine the developmental biology of *E. tabacivora*, aiming to make further progress in using the parasitoid in whitefly-control programs. This research might be particularly helpful in designing mass rearing protocols for using the parasitoid in inoculative biocontrol programs.

## 2 Materials and Methods

*Encarsia tabacivora* was collected in the Cañete valley, Peru from parasitized whitefly nymphs infesting sweet potato, and cassava. The rearing was carried out at the International Potato Center (CIP) in its host *B. tabaci* MEAN 1. Its unambiguous identification was carried out by Dr. Gregory Evans (USDA/APHIS). Voucher specimens (accession number: CIP-698-15) were kept in CIP’s entomology museum, Lima, Peru.

### 2.1 Insect rearing

### 2.1.1 Host

A colony of *B. tabaci* MEAN 1 was initiated with puparia collected from sweet potato in La Molina, Lima, Peru. The colony was maintained and mass-reared on sweet potato (cv. Costanero) in insect-proof cages (76.5 × 41 × 52 cm), which contained two sleeves for *in situ* manipulation, at controlled temperature (23-25°C), 70-75% relative humidity and photoperiod 12: 12 h light (L): dark (D).

### 2.1.2 Parasitoid

The rearing of *E. tabacivora* was initiated from parasitized whiteflies obtained from field-collected sweet potato and cassava leaves. Leaves were taken to the laboratory and immediately placed in acrylic plastic breeding boxes (32 × 32 × 30 cm). In the following days, emerging *E. tabacivora* adults were recovered using small glass tubes for easy transfer and release into rearing cages that contained sweet potato plants infested with nymphs of *B. tabaci* MEAN1.

Strips of parafilm dipped into a honey solution (2:1 honey: water) were placed inside the cages for providing a complementary food source. Adult wasps of the first generation were maintained in bioclimatic chambers at controlled temperature of 25°C. Wasps of the third and of following generations were reared at a temperature of 20°C.

For further rearing, six acrylic plastic cages with two sleeves were used (32 × 32 × 30 cm) that could hold 10 large vials (15 mm diameter × 45 mm long) containing sweet potato leaves infested with whitefly nymphs of the 3^rd^ and 4^th^ instar. Adult *E. tabacivora* wasps collected from the initial rearing were released into these cages. Every 10-15 days, the vials containing whitefly-infested leaves were renewed for providing further parasitation medium and avoid hyperparasitism. As previously, honey was provided in solution as additional food source for adult wasps.

### 2.1.3 Life cycle experiments

In life cycle experiments, 10 parasitoids were released into six polypropylene cylinders (12 × 9 ×12.5 cm) containing *B. tabaci* MEAN 1-infested sweet potato leaves (cv. Costanero). After a period of 6 hrs provided for parasitation, the parasitoids were removed using an aspirator, and parasitized nymphs were identified through the existence of oviposition punctures using a stereomicroscope. The position of parasitized nymph was marked on the leaves using a marker pen. Thereafter, the cylinders were stored at controlled temperature of 20°C and 70% RH.

Daily, 30 parasitized nymphs were collected from the cylinder for dissection to observe all developmental stages of the parasitoid until it had reached the adult stage. Dissections were carried out using a microscope (magnitude: 100×) in a drop of acid fuchsin diluted in distilled water (ratio 3:1). The most representative samples for each developmental instar were mounted on a slide and covered with balsam of Canada (Merck, Germany), following the preservation technique proposed by Noyes (1982). The development time for each development stage (2^nd^ and 3^rd^ larval instars, prepupae or black pupae) was determined. All stages of parasitoid development were photographed using an OLYMPUS (Model SZX-7) stereomicroscope equipped with an image analyzer. The images were used to measure the body size of the wasps using the ImageJ program (http://rsb.info.nih.gov/ij) by taking the following measurements: length and width of the body, width of the head, and length of the tail (the latter only in 1^st^ larval instars). In total, 570 nymphs were dissected.

## 3 Results

### 3.1 Biological and morphological characteristics of immature stages

The mean development time from egg to adult of *E. tabacivora* at a temperature of 20°C was 19.3 (SE ±0.17) days for females. The development stages were egg, three larval instars, prepupa and pupa (Fig. 1).

**Figure 1.**
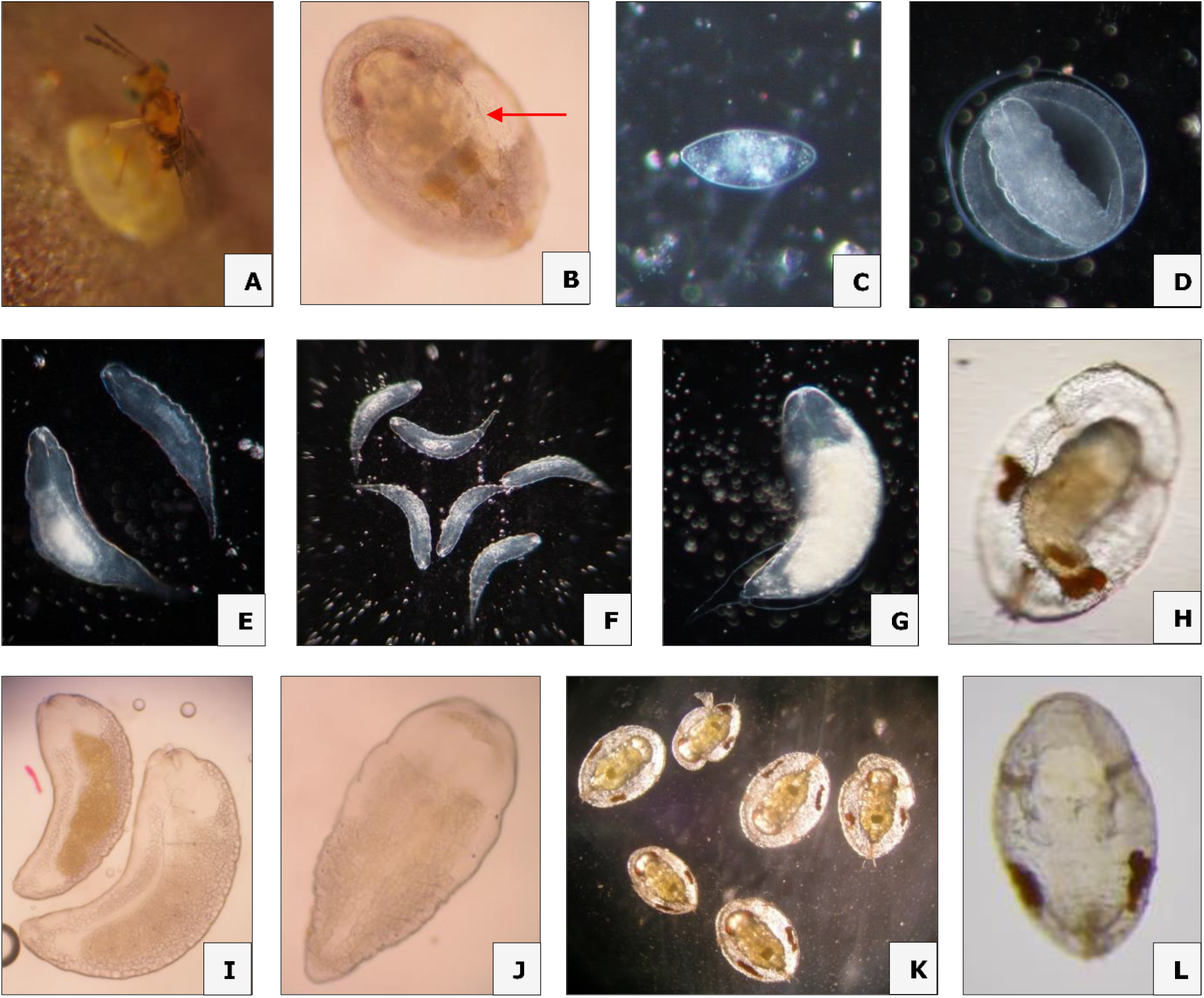
Observations in the development of *Encarsia tabacivora* females: (A) Female laying egg; (B) oviposition perforation; (C) freshly laid (elongated) egg; (D) developing egg showing formation of the first larval instar and the membrane with three layers, arrow - extra-embryonic membrane; (E) first larval instar with the presence of two larvae in a host; (F) first larval instar with the presence of six larvae in a host, showing superparasitism; (G) Second larval instar; (H) third larval instar leaving meconium, arrow: meconium; (I) differences between the second and third larval instar, blowhole arrow in the third instar larva; (J) prepupa, arrow: blowhole; (K) developing pupae; exuviates the host showing the exit orifice of the parasitoid.

When a female of *E. tabacivora* was exposed to nymphs of *B. tabaci* MEAN1, it began to move on the leaf palpating with the antennae; at times it quickly flapped its wings and cleaned its hind legs. When the female was attracted to a nymph, the female proceeded to examine the nymphs carefully, circling around and after properly choosing a nymph to parasitize, the female positioned itself over the nymph arching its abdomen in order to insert its ovipositor on the back of the cuticle (Fig. 1A). In some of these oviposition events the female remained up to 15 seconds with its ovipositor inserted in the host nymph; during this time the antennae was inclined forward. After oviposition, the female turned around and began to feed on the liquids that came out of the nymph, or she cleaned herself and walked around touching other leaves with the antenna in search of other host nymphs. There were also situations where the times were shorter than those mentioned above, so this was only an oviposition test, since the longer it lasted with the inserted ovipositor, the female was ovipositing as she remained immobile with the antennae tilted forward. Oviposition perforations appeared first light brown, but darkened later (Fig. 1B, puncture indicated by a red arrow). In some host nymphs, several oviposition punctures could be observed, indicating that more than one egg was likely laid in an individual host. Several 1^st^ instar larvae could be observed in the same host (Fig. 1E and F); however, only one individual 2^nd^ instar larva was always found in a single parasitized host.

The newly laid eggs were initially elongated (Fig. 1C and 2A) but changed to an elongated-oval shape over a few hours (Fig. 2B). The initially homogeneous egg content differentiated within a few hours, whereby the membrane layers separated, while the embryo was formed; as the size of the embryo increased, the shape of the egg rounded off. The color of the egg was translucent and towards the end of the egg’s development, the process of the 1^st^ larval instar formation could be observed (Fig. 2C and 1D). Egg development to the 1^st^ larval instar occurred in about 5-6 days; the body sizes during premature development [initial egg, the intermediate egg, and shortly before hatching (5-day old)] are shown in Table 1.

**Figure 2.**
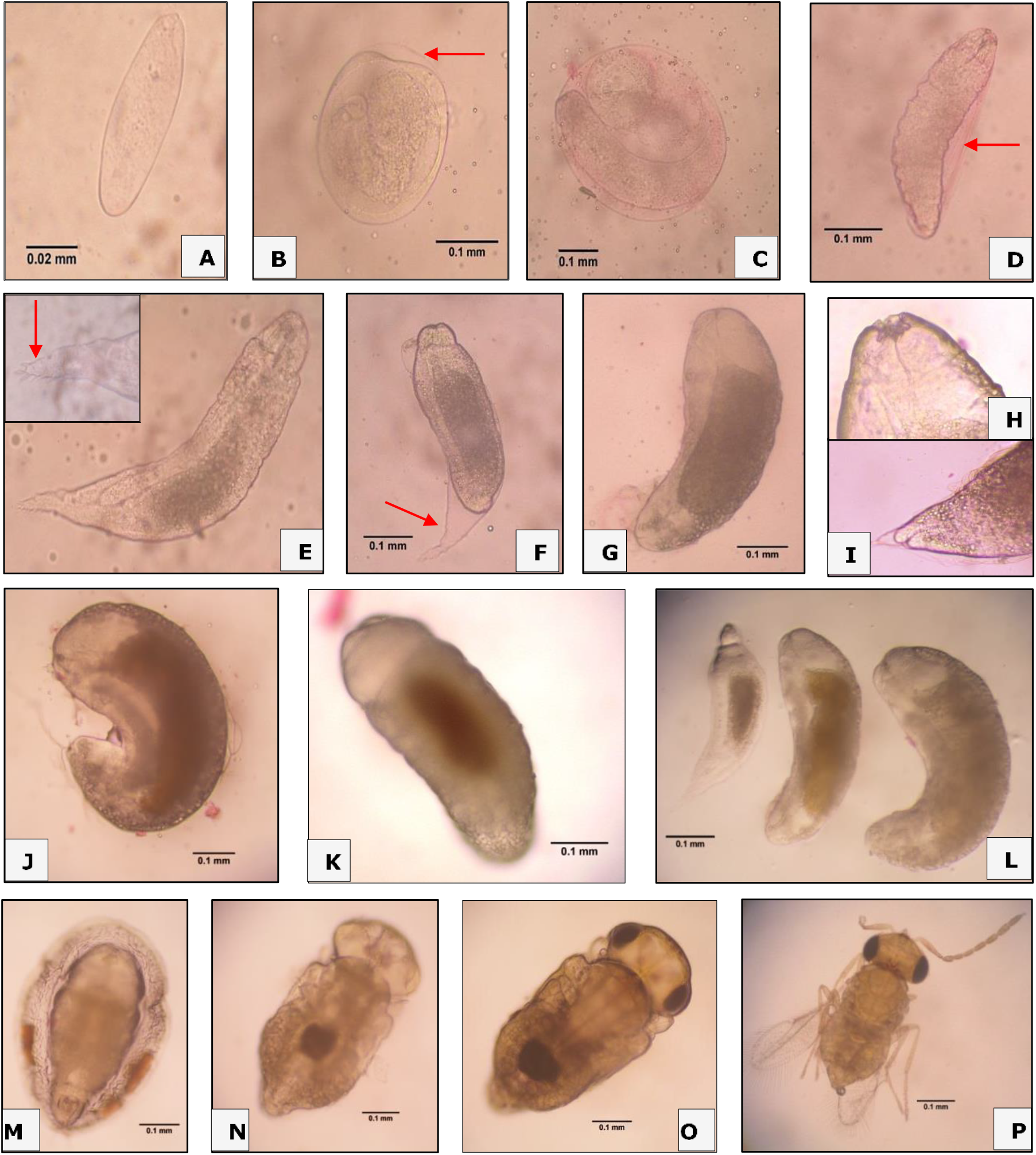
Development of immature stages and instars of *Encarsia tabacivora* females: (A) 1-day old egg; (B) developing egg with evidence of separation of the chorion from the embryo (arrow); (C) egg with larval formation process; (D) 1^st^ larval instar, arrow: remnants of the egg shell; (E) advanced 1^st^ larval instar, arrow: elongated tail with presence of setae in the terminal part; (F) 2^nd^ larval instar, arrow: exuvia of 1^st^ instar larvae; (G) 2^nd^ larval instar showing the body more enlarged and the size of the cephalic capsule increases, (H) view of the cephalic capsule and buccal apparatus; (I) View of the caudal appendix; (J) 3^rd^ larval instar; (K) third larval instar close to discharge of meconium; (L) comparative of shape and size between the 3 larval instar; prepupa, arrow: meconium; (N) unmelanized pupa; (O) pupa with advanced development; (P) formed adult.

Three larval instars were clearly differentiated by their morphology and size (Fig. 2, Table 1). Parasitoids of the 1^st^ instar have an elongated and transparent body (Fig. 2D and C). Neither the segments nor the respiratory spiracles were clearly visible during the early larval development. The head capsule was observed to be triangular; all contours had a smooth structure without setae or bumps (see head size in Table 1). The mandibles, which were in constant motion, were shaped like a “C”. The larvae had a thin, slightly elongated caudal appendage provided with setae (Fig. 2E, Table 1). The development time from the 1^st^ to the 2^nd^ instar was two days. The larvae were in constant motion in the host’s body fluid, but shortly before molting, their mobility decreased with increasing body size.

The 2^nd^ instar was legless and elongated, about 0.6-times thicker and wider than the 1^st^ instar and semi-arched in a ventral position (Table 1, Fig. 1). The early 2^nd^ instar had remnants of the anterior molt still attached to the last abdominal segments (Fig. 2F and 1G). The body surface was smooth and the body whitish, with the food ingested being visible through the yellow-brown color throughout the body (Fig. 2G). The spiracles were still not distinguishable, but parts of the tracheal system were apparent. The cephalic capsule increased in size (Table 1, Fig. 2H), and the buccal apparatus was perceptible; the jaws and digestive system was in continuous motion, sucking the contents of its host. The caudal appendix was markedly reduced and thus its mobility (Fig. 2I). The period of development from the 1^st^ to the 2^nd^ instar lasted about two days.

The 3^rd^ instar is very similar to the previous instars, but more arched in the ventral position, and with a body length of >0.5 mm and width of >0.2 mm it was larger than the previous instars (Fig. 2J). Other characteristics were similar; the body remained smooth, creamy white with yellow-brownish intestinal contents (Fig. 2K). The fully developed tracheal system showed nine pairs of spiracles; the first two are in the thoracic segments and the remaining seven in the abdominal segments, of which the second pair of abdominal spiracles was not fully developed (Figure 2L and 1I). The 3^rd^ instar completed its development within two days. The larva had consumed all the content of its host’s body and was visible by eye through the cuticle of the *B. tabaci* nymph. At the end of the 3^rd^ instar period, the parasitoid always acquired the (so-called) light form, that is, same position as the host, and discharged the meconium (small dark brown granules) along the inner margin of the empty pupa of the host (Fig. 1H).

The prepupae were creamy white, the internal organs were no longer visible through the cuticle, but the spiracles became noticeable (Fig. 1J). The prepupae were immobile remaining in the same location as the nymph of *B. tabaci* (light form, see body size in Table 1). Groups of meconium granules were observed on each side (Fig. 2M). Prepupae completed their development in less than 24 hrs.

The pupa was of the exarate type, the puparium envelope was transparent, showing the process of development, melanization and sclerotization (see body size in Table 1). At the beginning, the pupa was soft and whitish in color, the head with reddish compound eyes, and the thorax and abdomen could be differentiated (Fig. 2N). As pupae advanced in their development, the cuticle darkened (turning dark yellow) showing dark brown lateral stripes on the abdomen. The head was rounded with an average width of about 0.25 mm and 3 ocelli in the superior frontal position. The compound eyes changed to dark brown in color, the complete mouthparts took the anterior ventral position, and the antennae are inserted in the anterior part dorsal and are folded to the stergum (Fig. 2O). The thorax was narrower than the head, the wings were inserted in the lateral position and developed folded to the tergum. The abdomen had nine segments and a dark mass that probably correspond to the digestive system could be observed. Sclerotization of the pupae was gradual, the cuticle was harder and more solid towards the end of this stage. The pupae completed their development in seven days. The pupae generally developed in the same position as the whitefly pupae; that is, the position is facing the host’s belly, but in some cases, it could take the reverse position (Fig. 1K). When the pupae were close to emergence, the new adult turned to a supine position (Fig. 3A). This initiated a period of increased activity with intermittent motion of antennae and legs (Fig. 2P). For emergence, an irregular exit hole was drilled in the upper antero-dorsal part of the host exuvia on the opposite side to the operculum and close to the “T” suture through which the adult of the wasp emerges (Fig. 1L, 3B). The emergence process began with the head sticking out first and then the front legs. The front legs began immediately to swing until the rest of their body was released. All pupae of *E. tabacivora* were sexed as females.

**Figure 3.**
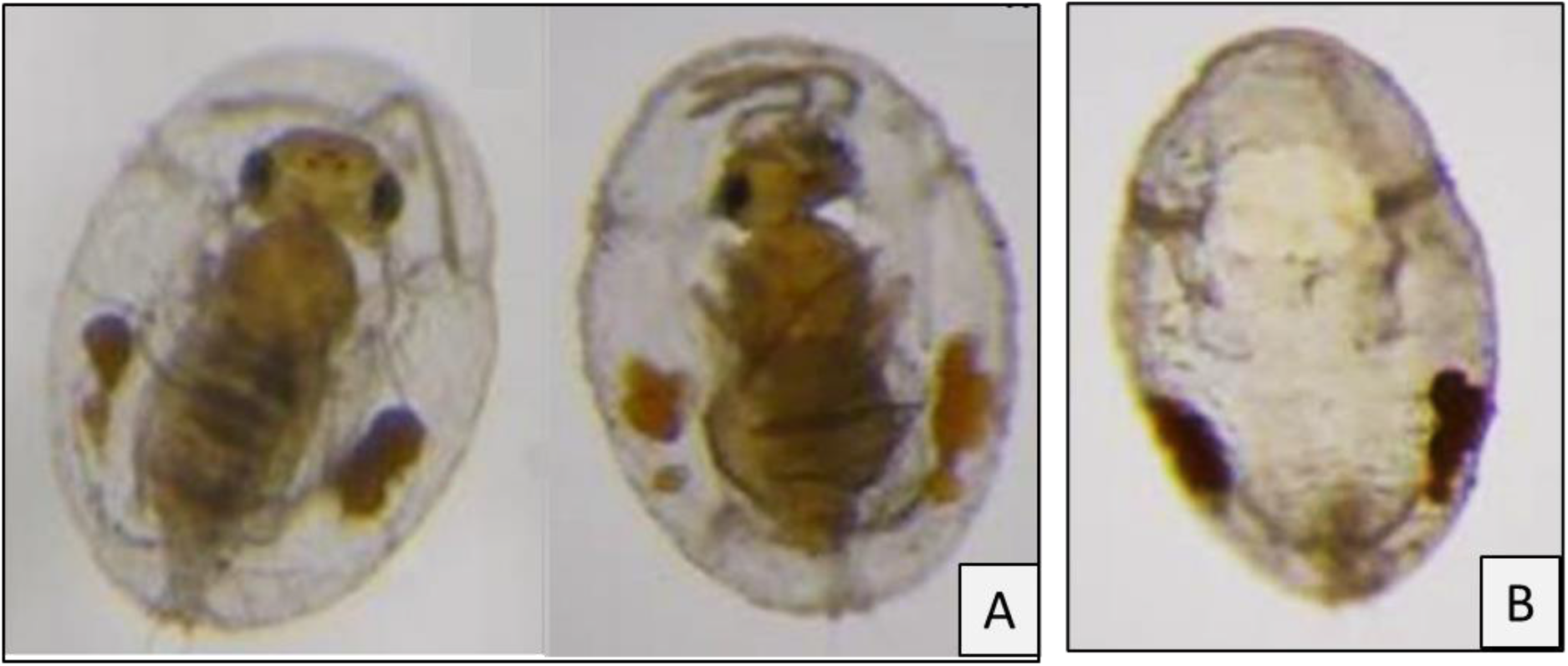
Emergence of *E. tabacivora* adults: (A) View of the location of the pupa about to emerge. (B) Exuvia of the host showing the exit orifice of the parasitoid.

### 3.2 Behavior and other characteristics of adults

#### 3.2.1 Mobility and flight capacity

Immediately after emergence, the adult cleaned its antennae with its front legs, its wings and abdomen using its hind legs (Fig. 4), walking slowly until its wings dried and were fully extended. The antennas were in constant motion. At temperatures of 20°C, the wasps showed the ability to fly very fast when they were threatened with a paintbrush and accelerated their movements by flying violently from one place to another.

**Figure 4.**
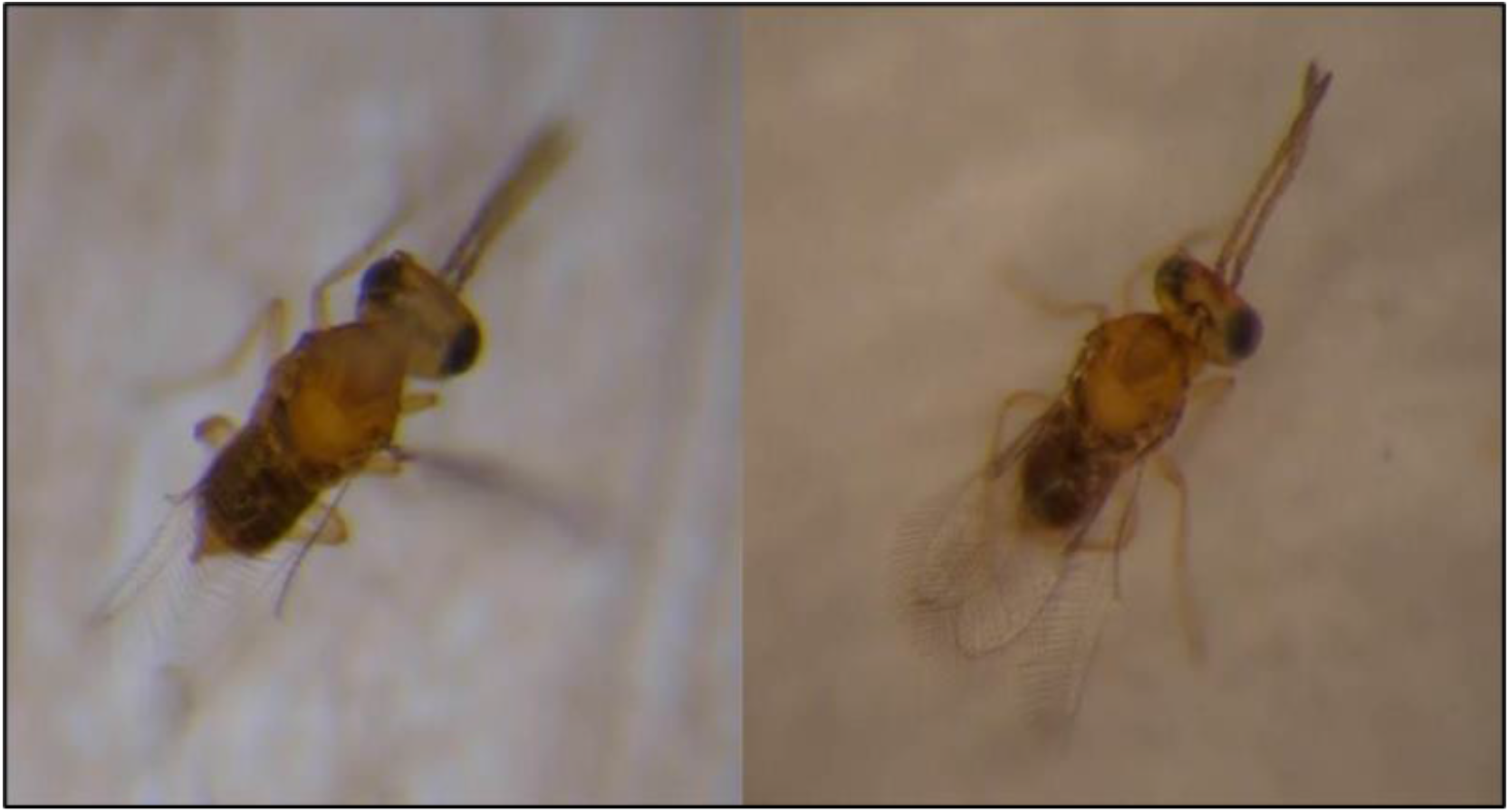
Newly emerged adults of *E. tabacivora*: Cleaning process of antennas and legs of newly emerged adults.

## 4 Discussion

*Encarsia tabacivora* has been reported in South America and the southern United States (Myartseva and Evans 2007). We have found that *E. tabacivora* prevails naturally in Peru and parasitizes nymphs of *B. tabaci* in cassava and sweet potato fields that are cultivated without the use of chemical pesticides. Agricultural fields regularly treated with chemical pesticides were parasitoid-free in our surveys. This suggests that the widespread use of chemical pesticides significantly deteriorates the abundance of *E. tabacivora* and other beneficial insects in these crops.

In this study, only *E. tabacivora* female adults emerged from parasitized whiteflies. The absence of males suggests that *E. tabacivora* exhibits thelytokous parthenogenesis like *E. formosa*. In *E. formosa*, asexual reproduction via thelytoky is induced by the infection with the bacterium Wolbachia (Zchori-Fein et al. 2001). The symbiotic bacterium is transmitted by the egg and restores the diploid state in eggs that have not been fertilized, producing females (Zchori-Fein et al. 1992). A strain of parthenogenesis-inducing endosymbiotic bacterium Cardinium has been reported to be associated with *E. tabacivora* populations in Brazil (Gebiola et al. 2017, Zchori-Fein et al. 2001, 2004). Without the infection with Wolbachia or Cardinium, parasitoids belonging to the order of Hymenoptera present arrhenotokous parthenogenesis, that is, the offspring of unfertilized females are composed only of males (De Bach 1985), which is the case of the species *Apsilophrys oeceticola* De Santis (Ávila and Redolfi 1990), *E. transvena* Timberlake (Antony et al. 2003), and *E. bimaculata* Heraty and Polaszek (Antony et al. 2004). In some species, bacterial symbionts cause cytoplasmic incompatibility, a form of reproductive incompatibility occurring when uninfected females mate with infected males, while the other three possible crosses between symbiont-infected and uninfected males and females produce normal numbers of offspring (Hunter et al. 2003, Perlman et al. 2006, Gebiola et al. 2015).

The development time of *E. tabacivora* from egg to adult emergence, determined at 20°C, was about 19 days, which is like the development time of 20.3 days reported for *Encarsia acaudaleyrodis* Hayat at the same temperature (Shishehbor and Zandi-Sohani 2011).

Newly laid eggs of *E. tabacivora* had an elongated shape which changed over the first hours to an elongated-oval shape until they became round before hatching, which is typical for many Encarsia species (Myartseva and Evans 2007). Our observations during day five of egg development of *E. tabacivora* referring to cell content variation and nucleic migrations with separation of membranous layers towards the peripheral regions are similar to the descriptions given by Antony et al. (2003) for the species *E. transvena* and Antony et al. (2004) for the species *E. bimaculata*.

In this study, more than one egg or 1^st^ instar larva of *E. tabacivora* were found in a host nymph of *B. tabaci*; however, only one individual adult emerged from one host. Similar observations were reported for *E. bimaculata* (Antony et al. 2004) and *Eretmocerus mundus* Mercet (Aphelinidae) (Gerling and Blackburn 2013). Van Alphen and Visser (1990) suggested that having two or more eggs in a host can increase the probability of producing an offspring compared to individual eggs, but Hagen and Van den Bosh (1968) stated that superparasitism only occurred under laboratory conditions. In contrast, Gerling and Foltyn (1987) reported that superparasitism occurred when host discrimination efficiency was reduced. Wylie (1983) found that parasitoid larvae took longer to develop in super parasitized hosts than in singly parasitized hosts.

The 1^st^ larval instar of *E. tabacivora* is elongated, transparent, showing a caudal appendage with protrusions at the ends like the species *E. pergandiella* (Gerling 1983, Antony et al. 2003), but different from *E. transvena* and *E. bimaculata* (Antony et al. 2003). In addition, *E. tabacivora* had a longer tail than *E. transvena* and *E. bimaculata* (Antony et al. 2003). The head of *E. tabacivora* is triangular with a notch containing a sickle shaped mandible like *E. transvena* and *E. pergandiella* (Antony et al. 2003).

The 2^nd^ larval instar of the genus Encarsia is generally quite similar to the 3^rd^ instar, but in endophagous species the airways are reduced. The 3^rd^ instar is very uniform and like other instars within the family Chalcidoidea (Myartseva and Evans 2007). In this study, morphological similarities were also found between the 2^nd^ and 3^rd^ instar, no remnants of molting from the previous stage were observed in the 3^rd^ instar, as observed by Antony et al. (2003, 2004) for *E. transvena* and *E. bimaculata*. For *E. tabacivora*, the determining observation to establish the change between larval instars was the presence of a long tail in the 1^st^ instar, the loss of this tail in the 2^nd^ instar, and the distinction of the nine pairs of respiratory spiracles clearly visible in the 3^rd^ instar.

The prepupa and pupa of *E. tabacivora* developed within nymphs of *B. tabaci* as do many species of Hymenoptera endoparasites (Myartseva and Evans 2007). The beginning of the prepupal stage was established after the 3^rd^ instar larva discharged the meconium granules on each side of the host and became immobilized. The melanization process in the pupa, which occurs in *E. tabacivora* and some other Encarsia species, made it possible to distinguish the formation of the various structures of the pupa. The reddish compound eyes, the black body with brown stripes on the abdomen, the formation of the antennas, as well as legs and wings could be clearly recognized through the transparent puparium cover, revealing its development. In *E. transvena* (Antony et al. 2003) and *E. bimaculata* (Antony et al. 2004), for example, this melanization does not occur until the end of its development, when the black cuticle enveloping the pupa is shed.

The body sizes of different development stages of *E. tabacivora* were almost 2-times larger compared to sizes determined for *E. bimaculata* (Antony et al. 2003) and *E. transvena* (Antony et al. 2004); however, some observed differences could be caused by the different temperature conditions with 20°C in our study and 25° to 30°C used in the study conducted by Antony et al. (2003, 2004), as well as by the applied dissection and assembly methodology. The body length of pupae of *E. tabacivora* was similar to *E. bimaculata* and *E. transvena*, but the body width of these two parasitoids was greater than of *E. tabacivora*.

Our study concludes that *E. tabacivora* is a naturally-occurring parasitoid of *B. tabaci* MEAN1 (Gennadius) in the Cañete valley of Peru. We recommend conducting additional field surveys to also confirm its wider distribution in other sweet potato production regions of Peru as well as to study its impact on its host by evaluating parasitism rates throughout the sweet potato production season. Life table studies at different constant temperatures would be very helpful to determine and understand the temperature-dependent growth potential of the parasitoid and for predicting and mapping potential release areas for the inoculative biological control of whiteflies. The knowledge gained on its development biology and described in this study can be used for establishing an effective mass-rearing system for this parasitoid. For this, we additionally recommended to identify the presence of symbiotic bacteria associated with *E. tabacivora* of Peru to confirm the cause of thelytokous parthenogenesis reproduction in this species.

## 5 Acknowledgements

We are thankful to Luis Goicochea, Jorge Peralta and Eva Huaman for supporting data collection. The research presented was undertaken as part of, and funded by, the CGIAR Research Program on Roots, Tubers and Bananas (RTB) and supported by CGIAR Fund Donors (https://www.cgiar.org/funders/). Funding from the Bill and Melinda Gates Foundation (grant OPP53344) to support open access publishing is gratefully acknowledged.

## Captions

Table 1.Development time and average body length and width measurements of immature stages of *Encarsia tabacivora* at constant temperature conditions of 20°C.

## Reference

Ávila T, Redolfi de Suiza I (1990) Biología de Apsilophrys oeceticola (Hymenopt.: Encyrtidae) parasitoide de Pebops sp. (Lepidopt: Cosmopterygidae). Rev Per Ent 33:113–118.

Antony B, Palaniswami MS, Henneberry TJ (2003) Encarsia transvena (Hymenoptera: Aphelinidae) development on different Bemisia tabaci Gennadius (Homoptera: Aleyrodidae) instars. Environ Entomol 32 (3):584–591. https://doi.org/10.1603/0046-225X-32.3.584

Antony B, Palaniswami MS, Kirk AA, Henneberry TJ (2004) Development of Encarsia bimaculata (Heraty and Polaszek) (Hymenoptera: Aphelinidae) in Bemisia tabaci (Gennadius) (Homoptera: Aleyrodidae) nymphs. Biol Control 30:546–555 https://doi.org/10.1016/j.biocontrol.2004.01.018

De Santis L (1981) Sobre dos especes de Encarsia (Hymenoptera, Aphelinidae) del Brasil parasitoides de Bemisia tabaci (Hymenoptera, Aleyrodidae) [On two species of Encarsia (Hymenoptera, Aphelinidae) from Brazil parasitoids of Bemisia tabaci (Hymenoptera, Aleyrodidae)]. Rev Bras Entomol 25:37–38

Díaz F, Ternero L (2002) Encarsia pergandiella parasitoide de Bemisia argentifolii en la región Chavimochic. Sociedad Entomológica del Perú. Resúmenes Convención Nacional de Entomología, Lima, Perú. p. 111.

Evans GA, Serra CA (2002) Parasitoids associated with whiteflies (Homoptera: Aleyrodidae) in Hispaniola and descriptions of two new species of Encarsia Förster (Hymenoptera: Aphelinidae). J Hymen Res 11(2):197–212

Gamarra H, Mujica N, Carhuapoma P, Kreuze J, Kroschel J (2016) Sweetpotato whitefly, Bemisia tabaci (Gennadius 1989) (Biotype B). In: Kroschel, J., Mujica, N., Carhuapoma, P., Sporleder, M. (eds.). Pest distribution and risk atlas for Africa. Potential global and regional distribution and abundance of agricultural and horticultural pests and associated biocontrol agents under current and future climates. International Potato Center (CIP), Lima, pp 85–99. ISBN 978-92-9060-476.1. DOI: 10.4160/9789290604761

Gerling D (1983) Observations of the biologies and interrelationships of parasites attacking the greenhouse whitefly, Trialeurodes vaporariorum (West), in Hawaii. Proc Hawaiian Entomol Soc 24:217–225. http://hdl.handle.net/10125/11153

Gerling D, Blackburn MB (2013) Immature development of Eretmocerus mundus (Hymenoptera: Aphelinidae). Arthropod Struct Dev 42:309–314. https://doi.org/10.1016/j.asd.2013.03.003

Gerling D, Foltyn S (1987) Development and host preference of Encarsia lutea (Masi) and interspecific host discrimination with Eretmocerus mundus (Mercet) (Hymenoptera: Aphelinidae) parasitoids of Bemisia tabaci (Gennadius), (Homoptera: Aphelinidae). J Appl Entomol 103:425–433. https://doi.org/10.1111/j.1439-0418.1987.tb01004.x

Gebiola M, Monti MM, Johnson RC, Woolley JB, Hunter MS, Giorgini M, Pedata PA (2017) A revision of the Encarsia pergandiella species complex (Hymenoptera: Aphelinidae) shows cryptic diversity in parasitoids of whitefly pests. Syst Entomol 42:31–59. https://doi.org/10.1111/syen.12187

Giorgini M, Monti M, Caprio E, Stouthamer R, Hunter M (2009) Feminization and the collapse of haplodiploidy in an asexual parasitoid wasp harboring the bacterial symbiont Cardinium. Heredity 102:365–371 https://doi.org/10.1038/hdy.2008.135

Hagen KS, van den Bosch R (1968) Impact of pathogens, parasites, and predators on aphids. Annu Rev Entomol 13:325–384. https://doi.org/10.1146/annurev.en.13.010168.001545

Hernández-Suárez E, Carnero A, Aguiar A, Prinsloo G, LaSalle J, Polaszek A (2003) Parasitoids of whiteflies (Hymenoptera: Aphelinidae, Eulophidae, Platygastridae; Hemiptera: Aleyrodidae) from the Macaronesian archipelagos of the Canary Islands, Madeira, and the Azores. Syst Biodivers 1:55–108 https://doi.org/10.1017/S1477200002001007

Hunter MS, Perlman SJ, Kelly SE, (2003) A bacterial symbiont in the Bacteroidetes induces cytoplasmic incompatibility in the parasitoid wasp Encarsia pergandiella. Proc R Soc London Ser B 270:2185–2190 https://doi.org/10.1098/rspb.2003.2475

Myartseva SN, Ruiz CE, Coronado BJM (2012) Aphelinidae (Hymenoptera: Chalcidoidea) de importancia agrícola en México. Revisión y claves. Departamento de fomento editorial de la Universidad Autónoma de Tamaulipas, 1st edn. Tamaulipas, México. ISBN: 978-607-7654-38-4.

Myartseva SN, Evans GA (2007) Genus Encarsia Förster of Mexico (Hymenoptera: Chalcidoidea: Aphelinidae) A revision, key and description of new species, Serie avispas parasitícas de plagas y otros insectos. Universidad Autónoma de Tamaulipas UAM Agronomia y Ciencias, Ciudad Victoria, Tamaulipas, México. ISBN 978-970-9031-37-9.

Noyes JS (2015) Universal Chalcidoidea Database. Natural History Museum. http://www.nhm.ac.uk/chalcidoids. Accessed 25 February 2015

Perlman SJ, Kelly SE, Zchori-Fein E, Hunter MS (2006) Cytoplasmic incompatibility and multiple symbiont infection in the ash whitefly parasitoid, Encarsia inaron. Biol Control 39:474–480. https://doi.org/10.1016/j.biocontrol.2006.05.015

Polaszek A, Manzari S, Quicke DLJ (2004) Morphological and molecular taxonomic analysis of the Encarsia meritoria species-complex (Hymenoptera, Aphelinidae), parasitoids of whitetlies (Hemiptera, Aleyrodidae) of economic importance. Zool Scr 33:403–421. https://doi.org/10.1111/j.0300-3256.2004.00161.x

Polaszek A, Evans GA, Bennett FD (1992) Encarsia parasitoids of Bemisia tabaci (Hymenoptera, Aphelinidae, Homoptera, Aleyrodidae): A preliminary guide to identification. Bull Entomol Res 82:375–392 https://doi.org/10.1017/S0007485300041171

Shishehbor P, Zandi-Sohani N (2011) Investigation on functional and numerical responses of Encarsia acaudaleyrodis parasitizing Bemisia tabaci on cucumber. Biocontrol Sci Technol 21(3):271–280. https://doi.org/10.1080/09583157.2010.543268

Supanta L (2017) La temperatura sobre la biología de Encarsia tabacivora Viggiani (Hym.: Aphelinidae) parasitoide de Bemisia tabaci (Gennadius) (Hem. Aleyrodidae). BSc Thesis,. Universidad Nacional Agraria La Molina. 125p.

Van Alphen J, Visser M (1990) Superparasitism as an adaptative strategy for insect parasitoids. Annu Rev Entomol 35:59–79 https://doi.org/10.1146/annurev.en.35.010190.000423

Viggiani G (1988) Le specie italiane del genere Encarsia Foerster. Boll Lab Entomol Agrar Filippo Silvestri 44:121–179. https://www.nhm.ac.uk/resources/research-curation/projects/chalcidoids/pdf_X/Viggia988d.pdf

Viggiani G (1985) Notes on a few Aphelinidae, with description of five new species of Encarsia Foerster (Hymenoptera: Chalcidoidea). Boll Lab Entomol Agrar Portici 42:81–94. https://www.nhm.ac.uk/resources/research-curation/projects/chalcidoids/pdf_X/Viggia985d.pdf

Wylie HG (1983) Delayed development of Microctomus vittatae (Hymenoptera: Braconidae) in superparasitized adults of Phyllotreta cruciferae (Coleoptera: Chrysomelidae). Can Entomol 115:441–422 https://doi.org/10.4039/Ent115441-4

Zchori-Fein E, Roush RT, Hunter MS (1992) Male production induced by antibiotic treatment in Encarsia formosa, an asexual species. Cell Mol Life Sci 48:102–105. DOI:10.1007/BF01923619

Zchori-Fein E, Gottlieb Y, Kelly SE, Brown JK, Wilson JM, Karr TL, Hunter MS (2001) A newly discovered bacterium associated with parthenogenesis and a change in host selection behavior in parasitoid wasps. Proc Natl Acad Sci USA 98:12555–12560. https://doi.org/10.1073/pnas.221467498

Zchori-Fein E, Perlman S, Kelly SE, Katzir N, Hunter MS (2004) Characterization of a ‘Bacteroidetes’ symbiont in Encarsia wasps (Hymenoptera: Aphelinidae): proposal of “Candidatus Cardinium hertigii”. Int J Syst Evol Microbiol 54:961–968. https://doi.org/10.1099/ijs.0.02957-0

