## Supplementary material for "Development of *Encarsia tabacivora* (Viggiani) (Hymenoptera: Aphelinidae) in nymphs of *Bemisia tabaci* (Gennadius) MEAN 1 (Hemiptera: Aleyrodidae)": Table 1

Table 1. Development time and average body length and width measurements of immature stages of *Encarsia tabacivora* at constant temperature conditions of 20°C.

| **Parasitoid status** | **n** | **Development time (days)** | **Lenght (mm ±STD)** | **Width (mm ±STD)** | **Head width (mm ±STD)** | **Tail length (mm ±STD)** |
|---|---|---|---|---|---|---|
| Starting egg | 31 | 2 | 0.091 ± 0.0037 | 0.043 ± 0.0021 | | |
| Intermediate egg | 17 | 1.5 | 0.157 ± 0.0040 | 0.107 ± 0.0037 | | |
| Advanced egg | 32 | 2 | 0.322 ± 0.0098 | 0.254 ± 0.0097 | | |
| 1$^{st}$ instar larva | 57 | 2 | 0.393 ± 0.0091 | 0.100 ± 0.003 | 0.055 ± 0.0013 | 0.045 ± 0.0021 |
| 2$^{nd}$ instar larva | 47 | 2 | 0.456 ± 0.0110 | 0.157 ± 0.0038 | 0.085 ± 0.0027 | |
| 3$^{rd}$ instar larva | 46 | 2 | 0.567 ± 0.0055 | 0.214 ± 0.0055 | 0.141 ± 0.0057 | |
| Prepupae | 20 | 1 | 0.577 ± 0.0252 | 0.253 ± 0.0103 | 0.224 ± 0.0099 | |
| Pupae | 39 | 7 | 0.595 ± 0.0025 | 0.255 ± 0.0061 | 0.248 ± 0.0033 | |
